# Chemically-informed Analyses of Metabolomics Mass Spectrometry Data with Qemistree

**DOI:** 10.1101/2020.05.04.077636

**Authors:** Anupriya Tripathi, Yoshiki Vázquez-Baeza, Julia M. Gauglitz, Mingxun Wang, Kai Dührkop, Mélissa Nothias-Esposito, Deepa D. Acharya, Madeleine Ernst, Justin J.J. van der Hooft, Qiyun Zhu, Daniel McDonald, Antonio Gonzalez, Jo Handelsman, Markus Fleischauer, Marcus Ludwig, Sebastian Böcker, Louis-Félix Nothias, Rob Knight, Pieter C. Dorrestein

## Abstract

Untargeted mass spectrometry is employed to detect small molecules in complex biospecimens, generating data that are difficult to interpret. We developed Qemistree, a data exploration strategy based on hierarchical organization of molecular fingerprints predicted from fragmentation spectra, represented in the context of sample metadata and chemical ontologies. By expressing molecular relationships as a tree, we can apply ecological tools, designed around the relatedness of DNA sequences, to study chemical composition.

## Main

Molecular networking^1^, introduced in 2012, was one of the first data organization approaches to visualize the relationships between fragmentation spectra for similar molecules from tandem mass spectrometry data in the context of metadata. It formed the basis for the web-based mass spectrometry infrastructure, Global Natural Products Social Molecular Networking^2^ (GNPS, https://gnps.ucsd.edu/) which sees ~200,000 new accessions per month. Molecular networking is used for a range of applications^3^ in drug discovery, environmental monitoring, medicine, and agriculture. While molecular networking is useful for visualizing closely related molecular families, the inference of chemical relationships at a dataset-wide level and in the context of diverse metadata requires complementary representation strategies. To address this need, we developed an approach that uses fragmentation trees^4^ and supervised machine learning^5^ to calculate all pairwise chemical relationships and visualizes it in the context of sample metadata and molecular annotations. We show that a chemical tree enables the application of various tree-based tools, originally developed for analyzing DNA sequencing data^6–9^, for exploring masss-pectrometry data.

We introduce Qemistree, pronounced *chemis-tree*, a software that constructs a chemical tree from fragmentation spectra based on predicted molecular fingerprints^10^. Molecular fingerprints are vectors where each position encodes a substructural property of the molecule. Recent methods allow us to predict molecular fingerprints from tandem mass spectra^11–15^. In Qemistree,we use SIRIUS^16^ and CSI:FingerID^13^ to obtain predicted molecular fingerprints. The users first perform feature detection^17,18^ to generate a list of observed ions, referred to as chemical features henceforth, to be analyzed by Qemistree (Fig. S1). SIRIUS then determines the molecular formula of each feature using the isotope and fragmentation patterns, and estimates the best fragmentation tree explaining the fragmentation spectrum. Subsequently, CSI:FingerID operates on the fragmentation trees using kernel support vector machines to predict molecular properties (2936 properties; Table S1). We use these molecular fingerprints to calculate pairwise distances between chemical features that are hierarchically clustered to generate a tree representing their structural relationships. Although alternative approaches to hierarchically cluster features based on cosine similarity of fragmentation spectra exist^19–21^, we use molecular fingerprints as it allows us to compare features based on a diverse range of structural properties predicted by CSI:FingerID. Additionally, as CSI:FingerID was shown to perform well for automatic *in silico* structural annotation^22^, we leverage it to search molecular structural databases to provide complementary insights into structures when no match is obtained against spectral libraries. Subsequently, we use ClassyFire^23^ to assign a 5-level chemical taxonomy (kingdom, superclass, class, subclass, and direct parent) to all molecules annotated via spectral library matching and *in silico* prediction.

Phylogenetic tools such as iTOL^24^ can be used to visualize Qemistree trees interactively in the context of sample information and feature annotations for easy data exploration. The outputs of Qemistree can also be plugged into other workflows in QIIME 2^25^ (many of which were originally developed for microbiome sequence analysis) or in R, Python etc. for system-wide metabolomic data analyses ^6,7,9, 26^. Qemistree is available to the microbiome community as a QIIME 2 plugin (https://github.com/biocore/q2-qemistree) and the metabolomics community as a workflow on GNPS^2^ (https://ccms-ucsd.github.io/GNPSDocumentation/qemistree/). The chemical tree from the GNPS workflow can be explored interactively (e.g. https://qemistree.ucsd.edu/).

To verify that molecular fingerprint-based trees correctly capture the chemical relationships between molecules, we generated an evaluation dataset with two human fecal samples, a tomato seedling sample, and a human serum sample. Mixtures of these samples were prepared by-combining them in gradually increasing proportions to generate a set of diverse but related metabolite profiles and untargeted tandem mass spectrometry was used to profile the chemical composition of these samples. Mass-spectrometry was performed twice using different chromatographic gradients causing a non-uniform retention time shift between the two runs. The data processing of these two experiments leads to the same molecules being detected as different chemical features in downstream analysis. In Figure 1a we highlight how these technical variations make the same samples appear chemically disjointed.

**Figure 1:**
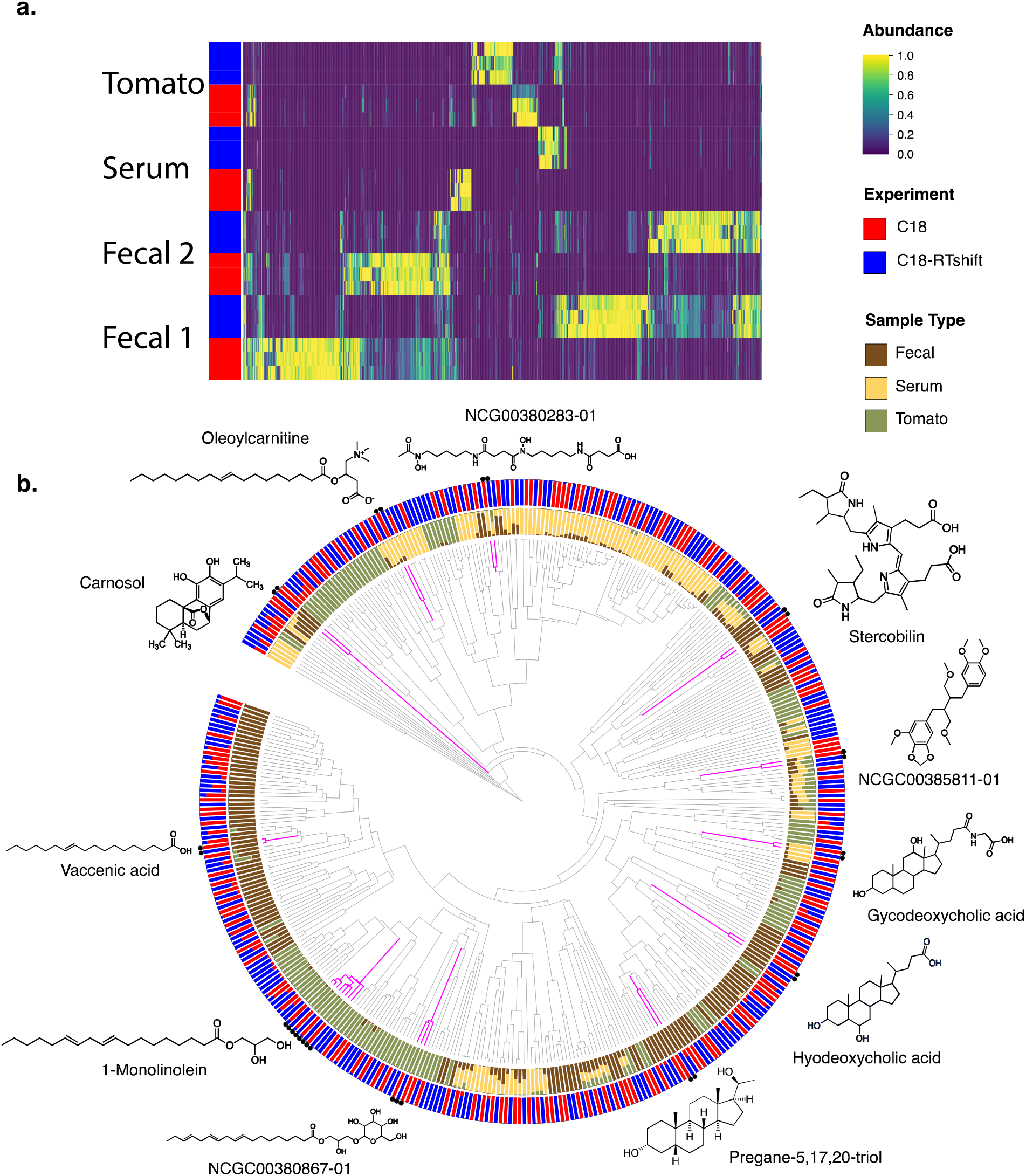
Qemistree mitigates aspects of technical artifacts by co-clustering structurally similar molecules across mass spectrometry runs. **a)** Sample (y-axis) by molecule (x-axis) heatmap of 2 fecal samples, tomato seedling samples, and serum samples in the evaluation dataset grouped by chromatography conditions. **b)** A chemical tree based on predicted molecular fingerprints representing the structural relationships between compounds detected in the evaluation dataset. Outer ring shows the relative abundance of molecules stratified by mass spectrometry run; inner ring shows the same stratified by fecal, serum and tomato samples in the evaluation dataset. Structurally similar molecules detected as different chemical features due to shift in retention time across mass spectrometry runs are clustered together; we highlight some examples of these artificially duplicated features around the tree. All structures shown are spectral reference library matches obtained from feature-based molecularnetworking^17^ in GNPS: (https://gnps.ucsd.edu/ProteoSAFe/status.jsp?task=efda476c72724b29a91693a108fa5a9d; Metabolomics Standard Initiative (MSI) level 3 annotation)^27^.

Using Qemistree, we map each of the spectra in the two chromatographic conditions (batches) to a molecular fingerprint, and organize these in a tree structure (Fig. 1b). Because molecular fingerprints are independent of retention time shifts, spectra are clustered based on their chemical similarity. This tree structure can be decorated using sample type descriptions, chromatographic conditions, and spectral library matches obtained from molecular networking in GNPS. Figure 1 shows that similar chemical features are detected exclusively in one of the two batches. However, based on the molecular fingerprints, these chemical features were arranged as neighboring tips in the tree regardless of the retention time shifts. This result shows how Qemistree can reconcile and facilitate the comparison of datasets acquired on different chromatographic gradients.

We demonstrate the use of a chemical hierarchy in performing chemically-informed comparisons of metabolomics profiles. In standard metabolomic statistical analyses, each molecule is assumed unrelated to the other molecules in the dataset. Some of the pitfalls of this assumption are highlighted in Figure 2a. Consider a scenario where we want to compare samples 1-3. An analysis schema that does not account for the chemical relationships among the molecules in these samples (Figure 2a, left), will assume that the sugars in samples 2 and 3 are as chemically related to the lipids in sample 1 as they are to each other. This would lead to the naive conclusion that samples 1 and 2, and samples 2 and 3 are equally distinct, yet they are not from a chemical perspective. On the other hand, if we account for the fact that sugar molecules are more chemically related to one another than they are to lipids, we can obtain a chemically-informed sample-to-sample comparison. Sedio and coworkers developed the chemical structural compositional similarity (CSCS) metric^28^ to account for relationships between molecules based on the similarity of their fragmentation spectra. While CSCS compares samples based on modified cosine scores obtained from molecular networking, we calculate chemical relationships based on structurally-informed molecular fingerprints. We express these relationships in the form of a hierarchy which enables the use of other tree-based tools for downstream data analyses. For example, in Figure 2a, we show that by using a tree of structural relationships between molecules in samples 1-3, we can apply UniFrac^9^, a tree-informed distance metric and demonstrate that the composition of sample 1 is distinct from samples 2 and 3.

**Figure 2:**
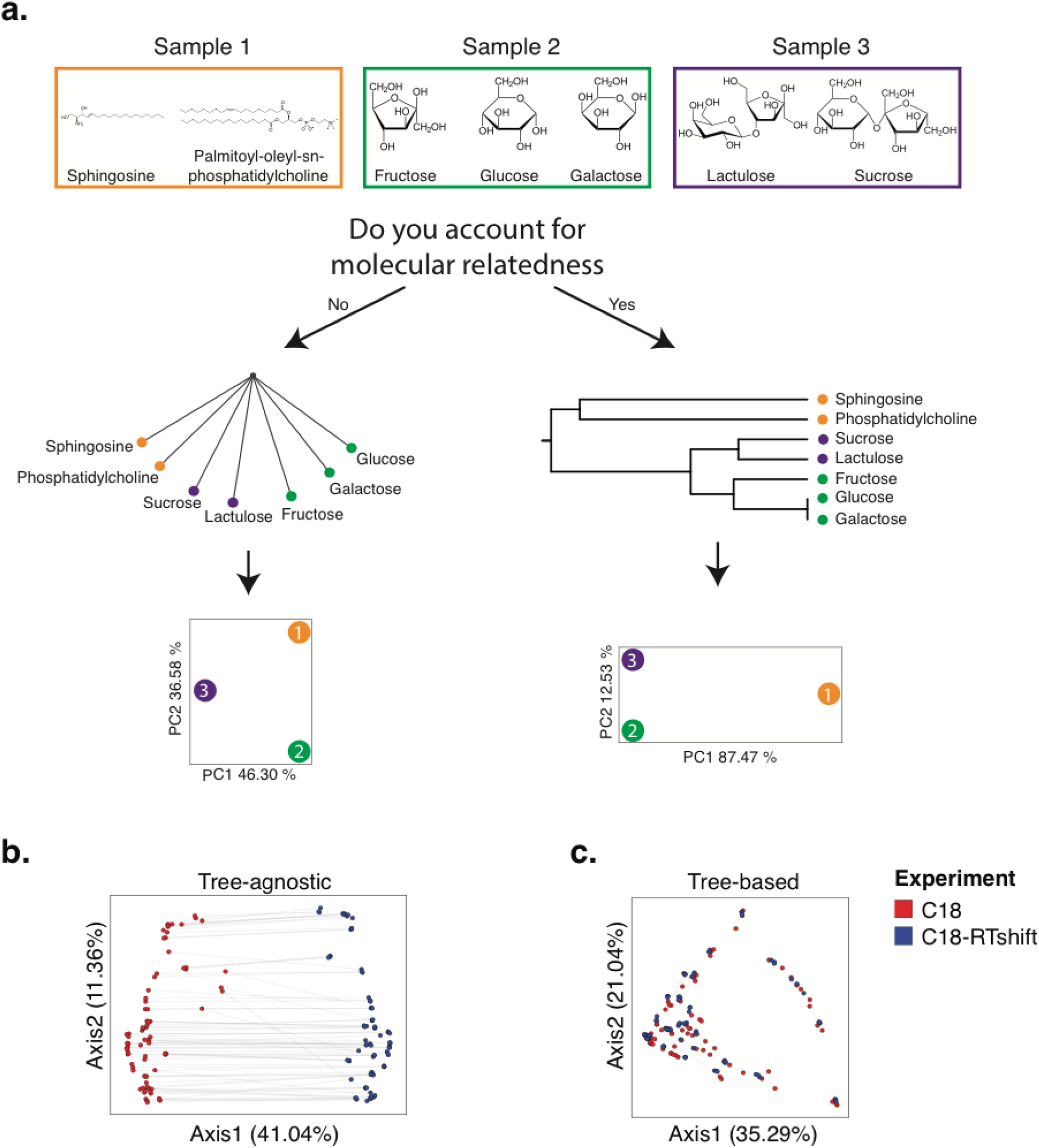
The pitfalls of assuming equal relatedness of molecules and the advantages of a chemical tree for sample comparison. **a)** A scenario where the goal is to compare the chemical composition in samples 1 (sphingosine and phosphatidylcholine), 2 (glucose, galactose, and fructose), and 3 (sucrose and lactulose). When we do not account for the chemical relationships between the molecules, i.e. assume that the lipid molecules in sample 1 are equally related to the sugars in samples 2 and 3 (left), we conclude that samples 1, 2, and 3 are similarly distinct. If we account for sugar molecules being more chemically related to one another than sugars are to lipid molecules(right), we can obtain a chemically-meaningful distance between samples. This is exemplified through a principal coordinates analysis (PCoA) of the computed UniFrac^9^ (tree-based) distances among samples; we see that samples 2 and 3 are more similar to each other, and sample 1 which is chemically distinct is separated along the primary axis of variation, when distances are computed using the chemical tree. **b, c)** PCoA of samples in the evaluation dataset colored by chromatography conditions. PCoA plot using tree-agnostic (Bray-Curtis^29^) distances which do not account for the chemical relationship between features detected across chromatography conditions (b) and tree-based (Weighted UniFrac^9^) distances which are based on the hierarchical relationships between molecules in the evaluation dataset (c).

The importance of comparing samples by accounting for their molecular relatedness is highlighted when we contrast the results from ignoring the tree structure (Fig. 2b) to those which integrate it (Fig. 2c). With the structural context provided by Qemistree, the differences between replicates across batches are comparable to the within-batch differences (Fig. S2). The retention time shift in this dataset leads to a strong technical signal that obscures the biological relationships among the samples (permutational ANOVA; tree agnostic^29^ pseudo-F=120.75, p=0.001 vs. tree informed^9^ pseudo-F=18.2239, p=0.001). We observed and remediated a similar pattern originating from plate-to-plate variation in a recently published study investigating the metabolome and microbiome of captive cheetahs^30^ (Fig. S3). In this study, placing the molecules in a tree using Qemistree reduced the observed technical variation (Fig. S3 a, c), and highlighted the dietary effect that was expected (Fig. S3 b, d). These results show how systematic and spurious molecular differences can be mitigated in an unsupervised manner using chemically-informed distance measures based on a tree structure.

As a case study, we used Qemistree to explore chemical diversity in a set of food samples collected as a part of the Global FoodOmics initiative (http://globalfoodomics.org). We selected a diverse range of food ingredients to represent animal, plant, and fungal groupings^31^. We first performed feature-based molecular networking using MZmine^17,18^ to obtain spectral library matches for a subset of the chemical features (~20% annotated with cosine cutoff > 0.7). Understanding the chemical relationships between different foods is challenging because most molecules within foods are unannotated. Using Qemistree, we collated GNPS spectral library matches and *in silico* predictions from CSI:FingerID to annotate ~91% of the chemical features (total 634 features after quality filtering) with molecular structures. Using ClassyFire^23^, we assigned a chemical taxonomy to 60% of these structures; the remaining 40% returned no ClassyFire taxonomy. Labeling annotations allowed us to retrieve subtrees of distinct chemical classes (Fig. 3a) such as flavonoids, alkaloids, phospholipids, acyl-carnitines, and O-glycosyl-compounds in food products. We propagated ClassyFire annotations of chemical features (tree tips) to each internal node of the tree and labeled the nodes by pie charts depicting the distribution in chemical superclasses (Fig. S4a) and classes (Fig. S4b) of its tips. The molecular fingerprint-based hierarchy of chemical features agreed well with ClassyFire taxonomy assignment, further demonstrating that molecular fingerprints can meaningfully capture structural relationships among molecules in a hierarchical manner. Furthermore, Qemistree coupled the chemical tree to sample metadata, revealing distinct chemical classes expected for each sample type. Branches representing acyl-carnitines were exclusively found in animal products (shades of blue; Fig. 3a). In contrast, honey, although categorized as an animal product, shared most of its chemical space with plant products, reflective of the plant nectar and pollen-based diet of honey bees. We observed a clade of flavonoids in both plant products and honey (Figs. 3a, S4b), but no other animal-based foods.

**Figure 3:**
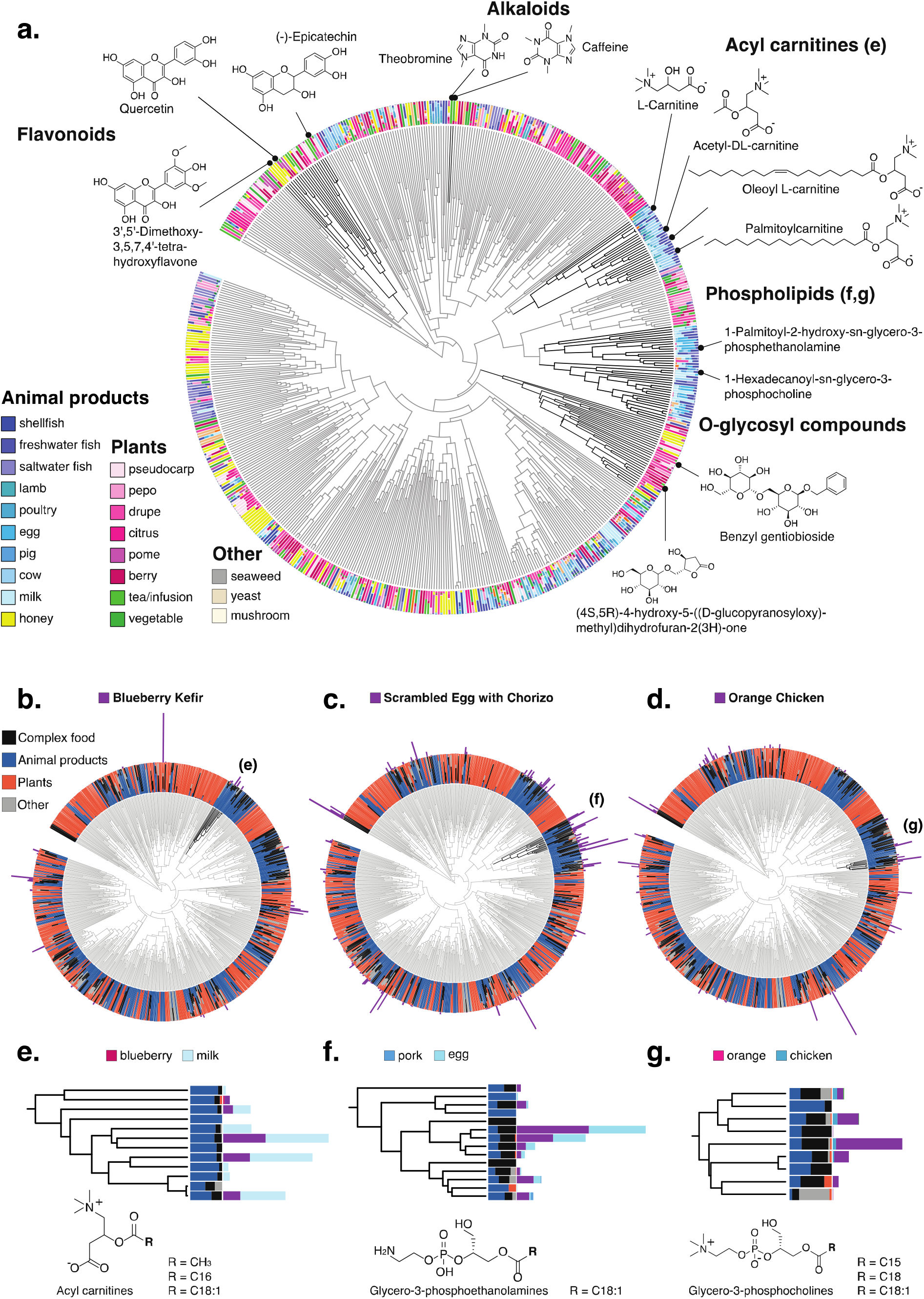
A chemical hierarchy of food-derived compounds based on predicted molecular fingerprints. **a)** A chemical tree based on molecular fingerprints representing the structural relationships between chemical features (tree tips) detected in food products (single ingredient i.e. simple foods; N=119). The tree is pruned to only keep tips that were assigned a structural annotation (SMILES) by either MS/MS spectral library match or *in silico* using CSI:FingerID. All structures shown are spectral reference library matches obtained from feature-based molecular networking in GNPS: (https://gnps.ucsd.edu/ProteoSAFe/status.jsp?task=ceb28a199d6b4f4fbf08490d9c96d631; MSI level 3 annotation^27^). The outer ring shows the relative abundance of each compound across a diverse range of food sources (panel a legend; parsed at ‘sample_type_group4’ of the Global FoodOmics Project ontology). We highlight clusters of compounds that are characteristic of specific food sources. For example, theobromine and caffeine are two closely related xanthine compounds (top center); they are primarily seen in teas (light green samples) and coffee beans (berry; purple). Similarly, acyl-carnitines and phospholipids (top right) are unique to different animal products (blues). We note that honey (highlighted in yellow), although annotated as an animal product, contains compounds that are primarily seen in plant sources (flavonoids, O-glycosyl compounds) and no other animal products. Flavonoids (top left) are observed in a range of fruit, vegetable, and honey samples (but no other animal products). **(b-d**) A hierarchy of the compounds observed in simple foods (above) and seven complex samples: two meals of orange chicken, a cooked cucumber and the sauce from a meal (schmorgurken), sour cream, blueberry kefir, and egg scramble with chorizo (N=126). The inner ring shows the relative abundance of each compound across simple animal products, plant products, fungi and algae (other) and the 7 complex foods (black). The absolute abundances of compounds in blueberry kefir (b), scrambled eggs with chorizo (c), and orange chicken (d) (outer bars) are overlaid on the tree to illustrate the shared and unique chemistry of complex foods. A compound subtree characteristic of each complex food in the tree is highlighted (black) and zoomed in **(e-g)**. (e) A subtree showing the absolute abundance of acyl carnitines in blueberry kefir and its primary ingredients (blueberry and milk). Similar subtrees showing phosphoethanolamine in scrambled eggs with chorizo (f), and phosphocholine in orange chicken (g).

While it is expected that a complex food such as blueberry kefir contains molecules from both blueberries and dairy, we can now visualize how individual ingredients and food preparation contribute to the chemical composition of complex foods. We noted that metabolite signatures that stem directly from particular ingredients, such as phosphoethanolamine from eggs, are present in egg scramble (Fig. 3c), but not in the other two foods highlighted (Fig. 3b and d). We can also observe the addition of ingredients in foods that were not listed as present in the initial set of ingredients. We were able to retrieve that there is black pepper in the egg scramble with chorizo and orange chicken, but that this signal is absent from the blueberry kefir (Fig. S5).

We show that our tree-based approach coherently captures chemical ontologies and relationships among molecules and samples in various publicly available datasets. Qemistree depends on representing chemical features as molecular fingerprints, and shares limitations with the underlying fingerprint prediction tool CSI:FingerID. For example, fingerprint prediction depends on the quality and coverage of MS/MS spectral databases available for training the predictive models, and these will improve as databases are enriched with more compound classes. Qemistree is also applicable in negative ionization mode; however, less molecular fingerprint scan be confidently predicted due to less publicly available reference spectra, resulting in less extensive trees.

In summary, we introduce a new tree-based approach for computing and representing chemical features detected in untargeted metabolomics studies. A hierarchy enables us to leverage existing tree-based tools, and can be augmented with structural and environmental annotations, greatly facilitating analysis and interpretation. We anticipate that Qemistree, as a data organization strategy, will be broadly applicable across fields that perform global chemical analysis, from medicine to environmental microbiology to food science, and well beyond the examples shown here.

## Data availability

The mass spectrometry data, metadata, and methods for the evaluation dataset have been deposited on the GNPS/MassIVE public repository^2,33^ under the accession number MSV000083306. The parameters used for molecular networking are available on GNPS: https://gnps.ucsd.edu/ProteoSAFe/status.jsp?task=efda476c72724b29a91693a108fa5a9d. The chemical hierarchy generated by Qemistree (version 2020.1.2) is available on iTOL^24^: https://itol.embl.de/tree/709513416494381587432576. The mass spectrometry data, metadata, and methods for Global Foodomics dataset have been deposited on the GNPS/MassIVE public repository^2,33^ under the accession number MSV000085226. The parameters used for molecular networking are available on GNPS: https://gnps.ucsd.edu/ProteoSAFe/status.jsp?task=ceb28a199d6b4f4fbf08490d9c96d631. The chemical hierarchy generated by Qemistree (version 2020.1.2) is available on iTOL^24^: https://itol.embl.de/tree/13711034118313741584046018.

## Code availability

All source code is publicly available under BSD-2-Clause on GitHub: https://github.com/biocore/q2-qemistree. Qemistree is also available as an advanced analysis workflow on GNPS: https://ccms-ucsd.github.io/GNPSDocumentation/qemistree/

## Acknowledgments

PCD was supported by the Gordon and Betty Moore Foundation (GBMF7622), the U.S. National Institutes of Health for the Center (P41 GM103484, R03 CA211211, R01 GM107550), and the University of Wisconsin-Madison OVCRGE; LFN was supported by the U.S. National Institutes of Health (R01 GM107550), and the European Union’s Horizon 2020 program (MSCA-GF, 704786). JJJvdH was supported by an ASDI eScience grant, ASDI.2017.030, from the Netherlands eScience Center—NLeSC. KD, MF, ML and SB were supported by Deutsche Forschungsgemeinschaft (BO 1910/20).

## Conflict of Interests

Mingxun Wang is a founder of Ometa Labs LLC. Pieter C. Dorrestein is a scientific advisor for Sirenas LLC. Kai Dührkop, Marcus Ludwig, Markus Fleischauer and Sebastian Böcker are founders of Bright Giant GmbH.

## Author contributions

PCD, AT conceived the concept and managed the project.

AT and YVB developed the algorithm and wrote the code for Qemistree.

AT and YVB contributed equally to the work.

LFN, RK, PCD supervised method implementation.

KD, MW, JJJvdH, ME, DM, and AG tested and provided suggestions on how to improve the method.

MW managed the deployment of Qemistree on GNPS.

AT and MW developed the GNPS-Qemistree Dashboard.

DA and AT wrote the documentation for the GNPS-Qemistree workflow.

YVB, QZ, and AT developed Qemistree-iTOL visualization.

LFN and MNE performed the mass-spectrometry for the evaluation dataset.

AT, YVB, and LFN analyzed and interpreted the evaluation data.

JMG performed mass spectrometry of the Global Foodomics samples.

AT, JMG analyzed and interpreted the Global Foodomics data.

KD, MF, ML, and SB supported the integration of SIRIUS, Zodiac, and CSI:FingerID.

PCD, AT, YVB, and RK wrote the manuscript.

LFN, JMG, MNE, JJJvdH, ME, KD, QZ, DM, AG, JH, MF, ML, SB, and RK improved the manuscript.

