## Supplementary Material for "Chemically-informed Analyses of Metabolomics Mass Spectrometry Data with Qemistree"

#### Supplemental Figures

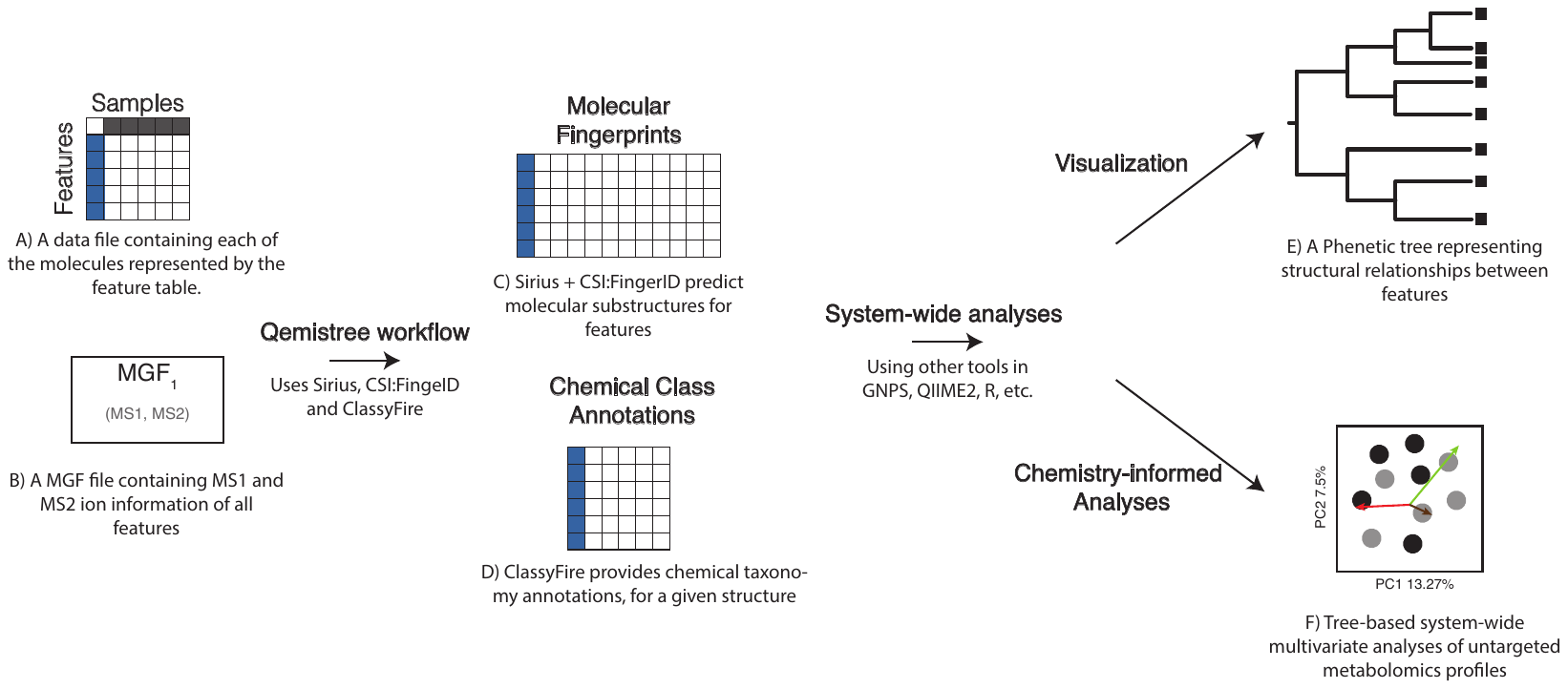

**Figure S1: End-to-end Qemistree analysis using GNPS and QIIME2.** Qemistree analysis can be performed using two required input files: 1) A table of molecule (or chemical feature) abundances per sample and 2) an MGF file with MS1 and MS2 ion information. These inputs can be generated by processing mass spectrometry files (.mzXML) through MZmine for feature detection. In Qemistree, these input files are processed through SIRIUS and CSI:FingerID to generate molecular fingerprints and *in silico* structural annotations (SMILES) per MS feature. We use the predicted molecular fingerprints to generate a phenetic tree of relationships between MS features based on sub-structural similarity. This tree can be visualized in iTOL for further data exploration. If the user inputs a sample metadata file, they can also visualize the abundances of each MS feature stratified by sample grouping of interest. Additionally, the qemistree queries ClassyFire to classify the structural annotations into chemical ‘kingdom’, ‘superclass’, ‘class’, ‘subclass’ and ‘direct parent’. We further allow the users to input a file with MS/MS spectral library matches (optional) into the workflow such that these library matches (typically, 2-20% of all MS features), instead of *in silico* annotation, are used for ClassyFire queries whenever available. All the outputs of the qemistree workflow can be analyzed further using QIIME 2 tools (such as tree-based alpha and beta diversity, mmvec^26^, songbird^32^) or explored in Python, R etc. as needed.

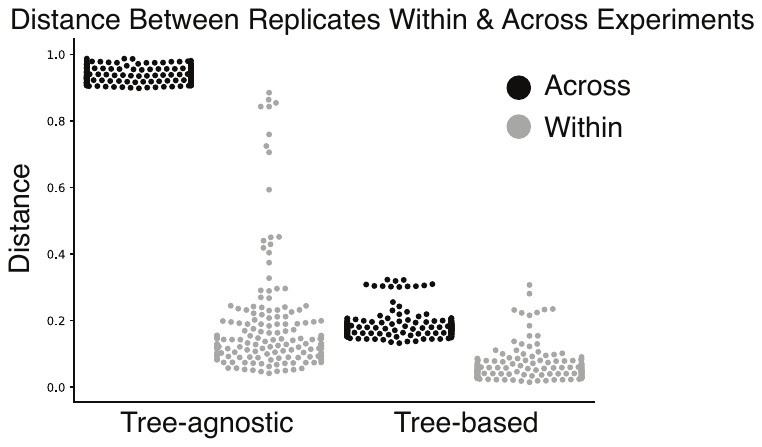

**Figure S2: Qemistree reduces the differences between biological replicates across mass-spectrometry runs**. A comparison of distances between sample replicates within and across chromatography gradients when using tree-agnostic (Bray-Curtis) distances and tree-based (Weighted UniFrac) distances.

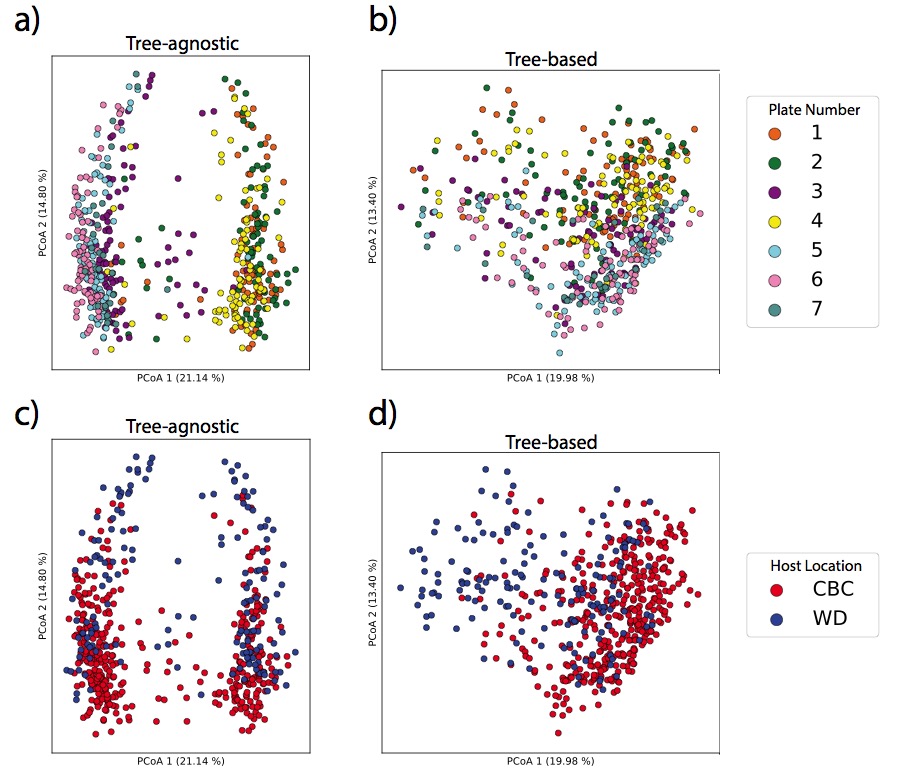

**Figure S3: Qemistree mitigates plate-to-plate variation in fecal metabolomics study to highlight a biologically-relevant effect.** a) Principal coordinate analysis (PCoA) of tree-agnostic distances (Bray-Curtis) colored by plate number (pseudo-F=32.39, p=0.001). b) PCoA of tree-informed distances (Weighted UniFrac) colored by plate number (pseudo-F=15.67, p=0.001). The same PCoA of (c) Bray-Curtis distances (pseudo-F=33.50, p=0.001) and (d) Weighted UniFrac distances (pseudo-F=48.42, p=0.001) colored by cheetah location which governed the diet of cheetahs. Data is available on the GNPS/MassIVE public repository^2,33^ accession number [MSV000082969](ftp://massive.ucsd.edu/MSV000082969/)**.**

CBC: Cheetah Breeding Center; WD: Wildlife Discoveries

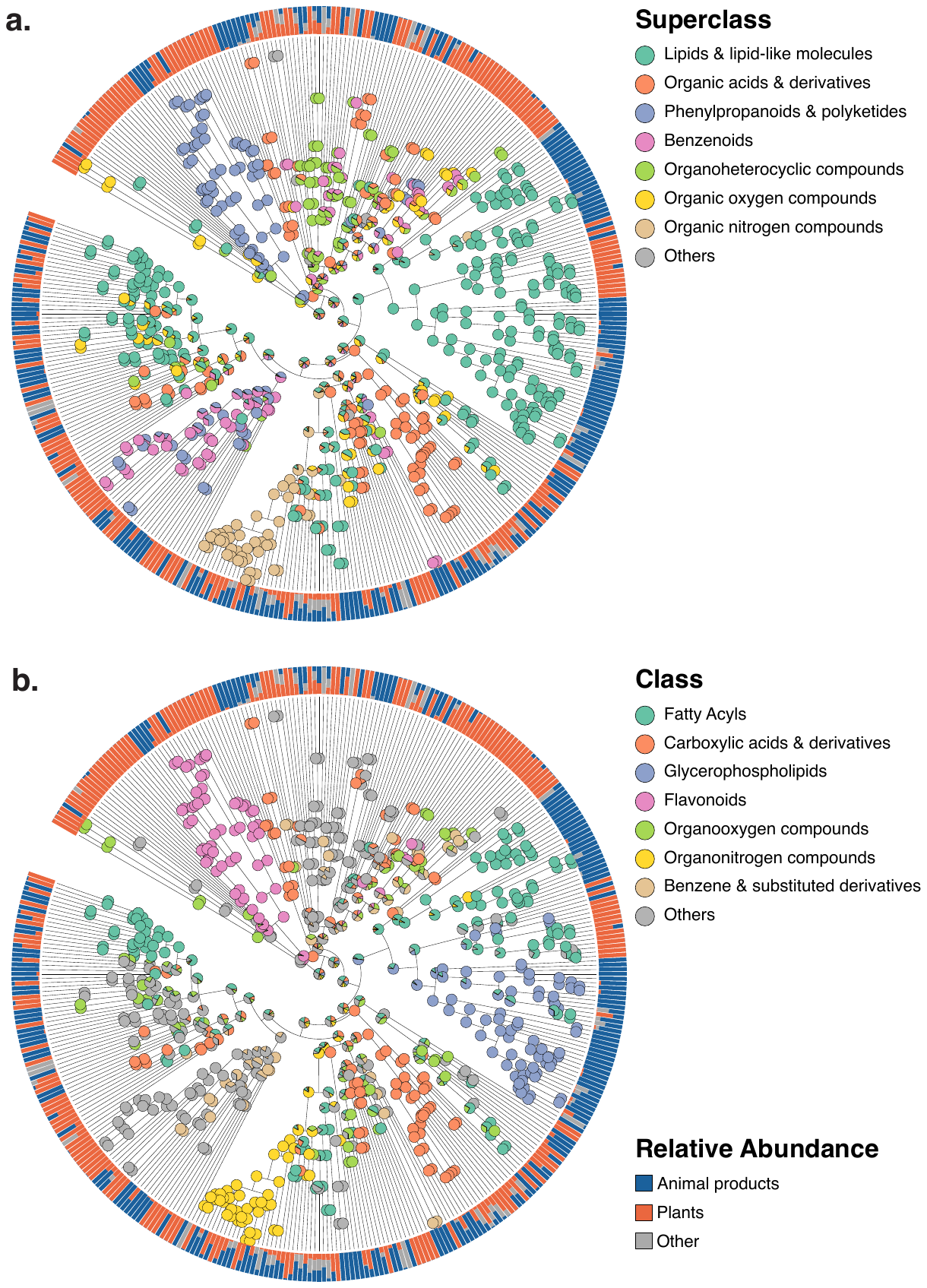

**Figure S4. Qemistree highlights chemical taxonomy of food-derived compounds.** Chemical hierarchy of compounds (tree tips) detected in simple food products (single ingredient foods, N=119). The tree is pruned to only tips that were assigned a chemical class using ClassyFire. Internal nodes are labeled by pie charts of the superclass **(a)** and class **(b)** level taxonomy of children tips. For instance, if a node has 10 children tips and 8 of them are assigned to class A and 2 to class B, then the pie chart will have two colors, with the angle being 8:2. Outer ring shows the relative abundance of each compound across simple animal products, plant products, and other (fungi and algae). The chemical hierarchy can be further explored using the following iTOL^24^ link: <https://itol.embl.de/tree/7095134164128581587333337>. For example, we observed two clades of chemical features classified as fatty acyls (b) such that the fatty acyl features found primarily in animal products are acyl carnitines (b; right) and the ones found in both plant and animal products are derivatives of linoleic acid (b; left).

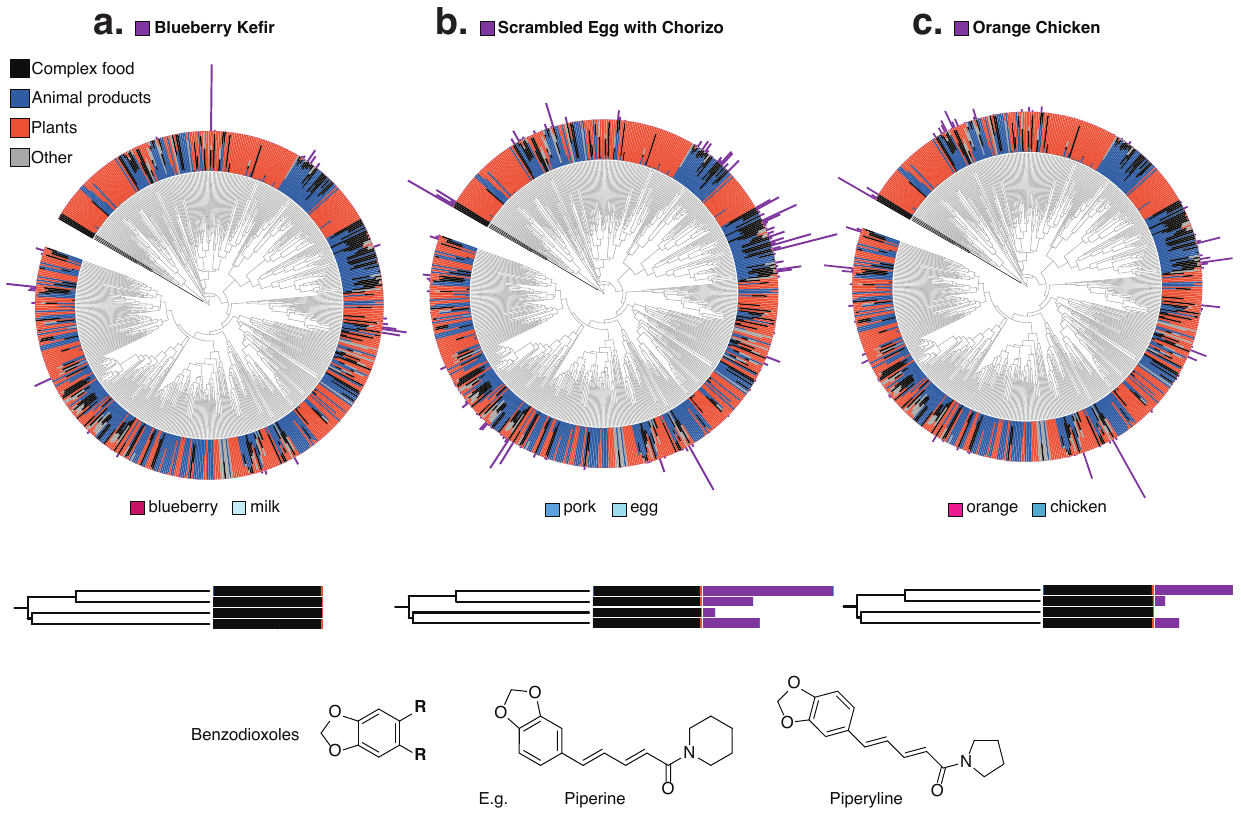

**Figure S5.**  **Chemical hierarchy of the compounds observed in simple foods (Figure 3a.) and seven complex samples**. **a,b,c)** 2 meals of orange chicken, a cooked cucumber and the sauce from a meal (schmorgurken), sour cream, blueberry kefir, and egg scramble with chorizo (N=126). Analogous to Figure 3b-d, the inner ring shows the relative abundance of each compound across simple animal products, plant products, fungi and algae (other) and complex foods. The absolute abundances of compounds in blueberry kefir **(a)**, scrambled eggs with chorizo **(b)**, and orange chicken **(c)** (outer bars) are overlaid on the tree to illustrate the shared and unique chemistry of complex foods. We highlight a classifier subtree annotated as benzodioxoles, compounds found in black pepper (in black) that are almost exclusively detected in complex foods. We overlay the absolute abundance of benzodioxoles in complex foods and their primary ingredients. These alkaloids are detected in scrambled eggs with chorizo (b) and orange chicken (c) but not in blueberry kefir (a) or the primary ingredients of these complex foods. This indicates that they are added during cooking, a likely assumption given the prevalence of black pepper in the western diet. The presence in an egg dish and meat dish coupled with the lack of signal in blueberry kefir also corresponds with the traditional use of this spice.

#### Online Methods

##### Qemistree algorithm

The Qemistree workflow uses MS1-based feature tables and MS1, MS2 fragment ion information (MGF file format) as inputs (Figure S1). These inputs can be generated by processing untargeted mass spectrometry data using MZmine^17^ following the Feature-Based Molecular Networking method^18^ (example batch file that can be used to perform feature detection and generate the inputs for Qemistree can be found here: [MSV000085226](ftp://massive.ucsd.edu/MSV000085226/)). The files exported from MZmine with the *Export/Submit to GNPS* and *SIRIUS Export* module, and are then imported into QIIME2^25^ as the following semantic types: FeatureTable[Frequency] (for the feature table) and MassSpectrometryFeatures (for the ion information).

```
# PREPROCESSING:
Use mzXML files from the instrument
Perform feature detection using MZMine2
Export sirius MGF and feature table (row m/z, row ID, feature area under the curve per sample)
Convert the feature table to FeatureTable[Frequency] for QIIME2
Create a FeatureData[Molecules] file for QIIME2 using ‘row ID’ and ‘row m/z’
Import the MGF file as MassSpectrometryFeatures for QIIME2
```

We use SIRIUS (version 4.0.1), ZODIAC^34^ and CSI:FingerID to predict molecular substructures within mass spectrometry features in the MGF files imported as MassSpectrometryFeatures. SIRIUS computes fragmentation trees for each molecular formula candidate of a feature (using PubChem database by default) and ranks these by score. SIRIUS uses MS1 spectrum in the MGF file to determine the candidate ion adduct(s) to be used for the fragmentation tree computation of each feature. ZODIAC takes the top SIRIUS candidates as input and re-ranks molecular formula candidates considering reciprocal compound similarities in the dataset to increase correct molecular formula assignments. Subsequently, CSI:FingerID predicts molecular fingerprints for each feature based on the molecular formula with the highest ZODIAC score.

Note that all spectra provided to the Qemistree pipeline do not necessarily produce a fingerprint. Indeed, SIRIUS does not compute fragmentation trees for multiply charged compounds and CSI:FingerID does not predict molecular fingerprints from spectra with less than 3 explained peaks. To ensure that high confidence molecular formulas are used in Qemistree, we only consider compounds with a ZODIAC score above 0.98 (described in ref. 34).

```
# SUBSTRUCTURE PREDICTION:
For each feature with MS2 spectra in the MGF file:
Compute fragmentation trees (using Sirius)
       Re-rank molecular formula candidates on the complete dataset (using Zodiac)
Predict fingerprints based on best molecular formula assignment (using CSI:FingerID)
```

A dataset *M* (i.e. a set of exports from MZmine) is a matrix of size *n* rows by *l* columns. Each row represents a molecule (*m_1_, m_2_, … m_n_*), and each column represents a molecular substructure feature. As such, each molecule *m_i_* is composed of a vector (with length *l*) of predicted probability values (one for each SIRIUS-generated molecular substructure). We remove from our analyses the features without a corresponding vector *m_i_*. In our tests, we have observed that for each dataset 10-15% of the input features are discarded.

For indexing purposes, we relabel each molecule *m_i_* with the MD5-checksum of the predicted fingerprint vector. The motivation to apply the MD5 hashing function is to assign a unique identifier to each feature, which is particularly useful when comparing datasets independently processed using Mzmine. If two distinct molecules (*i*, *j*) have identical checksums i.e. *md5(m_i_) = md5(m_j_)*, then we aggregate those two vectors such that all rows in *M* are unique. This operation is also propagated down to the table of molecular intensities, in that context intensities are added together.

To co-analyze multiple datasets *M_1_, M_2_, … M_k_* , we combine the matrices into a new dataset *M^*^*. For any two repeated molecules *m_i_* and *m_j_* in *M^*^* we merge their intensities and values as described before. Lastly, we create a hierarchy of chemical relationships *T* using a distance matrix *D* measuring the distance between all pairs of molecules in *M^*^*. For qualitative substructure comparisons, we use the Jaccard distance metric and a threshold of 0.5. Otherwise, we use the Euclidean distance with the original probability vectors. With *D*, we cluster the molecules in a hierarchical fashion using the unweighted pair group method with arithmetic mean (UPGMA). The tips in the resulting tree *T* have a one-to-one correspondence with all the molecules *m_i_* in *M^*^*.

```
# HIERARCHY CREATION (meta-analysis)
For each fingerprint, feature table in DATASETS:
Collate fingerprints into a matrix of features by fingerprints
Match the tuple to have the exact same features and same order
Merge all the fingerprints and feature tables
(use MD5 hash  of fingerprint vectors to merge identical fingerprints)
Compute a distance matrix between features using fingerprints (quantitatively or qualitatively)
Build a hierarchical tree based on the distance matrix
```

##### Evaluation dataset

**Sample preparation and extraction.** Four samples were used in the gradient benchmarking dataset: 1) the *“serum*” sample consists of the NIST SRM 1950 reference sample made of human serum spiked with compounds ^35^ 2) Two human fecal samples from the American Gut Project^36^ obtained from a single male individual with a 35 days interval (Sample *fecal-1* “ 11-10-2013, and *fecal-2 :* 12-14-2013), and 3) the “*tomato*” seedling sample (*Solanum lycopersicum* plant) was prepared using 3 weeks post-germination specimens (fresh whole seedlings were used). The NIST SRM 1950 sample (1mL), two fecal samples (210 mg of fresh material each), and the tomato seedlings (800 mg of fresh material) were dissolved in 1 mL of 7/3 methanol/water in a 1 mL polypropylene round-bottom tube (QIAGEN), and homogenized in a tissue-lyser (Tissue Lyser II, QIAGEN) at 25 Hz for 5 min. The tubes were then centrifuged at 15,000 rpm for 15 min, and 600 µL of the supernatant was collected and loaded on solid-phase extraction cartridges (Oasis HLB, Waters) made of hydrophilic-lipophilic balance stationary phase (30 mg and 30 µm particle size), that were first activated with 100% methanol, and 100% water (1mL each). After loading the supernatants on the cartridges, washing elution was carried out with 95/5 methanol/water (1 mL), and the samples were eluted with 7/3 methanol/water (2mL), followed by 100% methanol (1mL). The samples were dried down with a vacuum concentrator (Centrivap, Labconco) and resuspended in 2.5 mL of 7/3 methanol/water containing 0.5 µM of amitriptyline as an internal standard. Samples were prepared by mixing the four different samples in various proportions. The resulting extracts were analyzed by mass spectrometry but also used to prepare mixtures of these samples in different ratios. For example, the *serum* and *tomato samples* were mixed in the following ratios: 100/0, 75/25, 50/50, 25/75, 0/100.

**Liquid chromatography and mass spectrometry experiments.** Samples were analyzed using ultra high performance liquid chromatography (Vanquish, Thermo Scientific) coupled to a quadrupole-Orbitrap mass spectrometer (Q Exactive, Thermo Scientific). The quadrupole-Orbitrap mass spectrometer (Q Exactive, Thermo Scientific) was fitted with an electrospray source (HESI-II) operating in positive ionisation mode. The source used the following parameters: spray voltage, +3500 V; heater temperature, 437.5°C; capillary temperature, 268.75°C; S-lens RF, 50 (arb. units); sheath gas flow rate, 52.5 (arb. units); and auxiliary gas flow rate, 13.75 (arb. units). The samples were acquired in non-targeted MS^2^ acquisition mode, with up to four MS^2^ scans of the most abundant ions per MS1 scan. The spectra were recorded from 0.48 to 17 min. The following parameters were used for full MS scan: resolution (35,000), Automatic Gain Control target (1.0 x 10^6^), maximum injection time (125 ms), scan range (150-1500 m/z). For the data-dependent in MS^2^, the following parameters were used: resolution (17,500), AGC target (2.5 x 10^5^), maximum injection time (125 ms), loop count (4), isolation window (1.5 m/z) fixed first mass (70 m/z). (70-1500 m/z) and up to four MS/MS scans of the most abundant ions per duty cycle. Higher-energy collision induced dissociation was performed with a normalized collision energy of 30 (20, 35, 50). The data-dependent settings were set as follows: minimum AGC (1.25x 10^4^ [intensity threshold 1.0 x 10^5^]), apex trigger 3 to 15 s, charge exclusion 3-8 and > 8, exclude isotopes (on), dynamic exclusion (14.0 s).

**Mass spectrometry data processing.** Thermo mass spectrometry data (.RAW) were converted to *m/z* extensible markup language (mzML)^37^ in centroid mode using MSConvert ProteoWizard^38^ (release 201812). The mzML files were processed with MZmine toolbox ^17^ (version 2.38) on Ubuntu 18.04 LTS 64-bits workstation (intel Xeon 5E-2637, 3.5 GHz, 8 cores, 64 Go of RAM) following the Feature-Based Molecular Networking method.^18^

##### Global FoodOmics dataset

**Sample preparation and extraction.** Samples were collected, extracted, and MS data were acquired as a part of the Global FoodOmics project according to the sampling and data acquisition protocols described in Gauglitz et al., 2020 Food Chemistry. Briefly, 126 food samples were selected from the Global FoodOmics dataset. 119 simple food samples (simple in contrast to complex and defined as a single-ingredient food) were selected to cover a broad spectrum of fruits, vegetables, meat and fungi. Each food was represented in at least triplicate in the data subset. Additionally 7 complex samples were selected that contained simple foods from the simple food subset in their ingredient lists. The complex foods were from two separate meals of orange chicken, a cooked cucumber and the sauce from a meal (schmorgurken; in a tomato and sour cream sauce), sour cream, blueberry kefir, and egg scramble with chorizo. Sample metadata describes the food samples based on a food hierarchy beginning with plant vs. animal vs. fungus (sample_type_group1) and increasing in detail down to persian cucumber vs. cherry tomato etc. (sample_type_group6)

Briefly, samples were extracted in 95% LC-MS grade Ethanol; 5% LC-MS grade water. Samples were analyzed using the same LC-MS/MS setup and software as described above for the maXis II QTOF mass spectrometer (Bruker Daltonics), using a Phenomenex Kinetex C18 1.7 µm (100A) 100 x 2.1 column equipped with a guard cartridge (Phenomenex). The instrument tuning and internal calibrant remained the same as described above. MS spectra were acquired in a positive ion mode in the range m/z 50–1,500. The mobile phases consisted of A (100% water + 0.1% formic acid) and B (100% acetonitrile + 0.1% formic acid), and the flow rate was set to 0.5 µL/min throughout the experiment, and the column maintained at 40℃.

**Mass spectrometry data processing.** The mass spectrometry data (.d) were converted to .mzXML with lock mass calibration applied using CompassXport batch mode in Data Analysis 4.4 software (Bruker Daltonics, Bremen, Germany) running on a Windows 10 PC. The mass spectrometry data was processed with MZmine toolbox ^17^ (version 2.38) using the parameters outlined in an XML batch file (see Data availability).

**Multivariate comparisons.** To evaluate the benefits of using a tree for multivariate analysis, we generated pairwise sample distances using Bray-Curtis^29^ (tree-agnostic) and Weighted UniFrac^9^ (tree-informed). Both of these metrics compare samples quantitatively i.e. based on the abundances of each feature. Notably, UniFrac weights the distances based on the shared branches of the tree used for computation.
